# Sleep Induction by Mechanosensory Stimulation in *Drosophila*

**DOI:** 10.1101/2020.04.14.041061

**Authors:** Arzu Öztürk-Çolak, Patrick D. McClanahan, Joseph R. Buchler, Sho Inami, Andri Cruz, Christopher Fang-Yen, Kyunghee Koh

## Abstract

People tend to fall asleep when gently rocked or vibrated. Experimental studies have shown rocking promotes sleep in humans and mice. The prevailing “synchronization” model proposes synchronization of brain activity to mechanosensory stimuli mediates the phenomenon. The alternative “habituation” model proposes habituation, a form of non-associative learning, mediates sleep induction by monotonous stimulation. Here we show that gentle vibration promotes sleep in *Drosophila* in part through habituation. Vibration-induced sleep (VIS) leads to the accrual of homeostatic sleep credit, is associated with reduced arousability, and can be suppressed by heightened arousal. Sleep induction improves over successive blocks of vibration and exhibits stimulus specificity, supporting the habituation model. Multiple mechanosensory organs mediate VIS, and the magnitude of sleep gain depends on the vibration frequency and genetic background. Our findings suggest habituation is a major contributor to VIS, but synchronization of brain activity may play a role under certain stimulus conditions.

## Introduction

Anecdotal observations suggest that babies sleep better when gently rocked or bounced and people tend to fall asleep during long car rides. Several experimental studies have confirmed that rocking promotes sleep in infants, adult humans, and mice (Bayer et al., 2011; Kompotis et al., 2019; Korner et al., 1978; Perrault et al., 2019). Yet the underlying mechanisms are not well understood.

A prevailing model of how sensory stimulation promotes sleep, which we will term the synchronization model, is that it enhances the synchronous neural activity and boosts sleep slow waves (Bellesi et al., 2014; Perrault et al., 2019). Transcranial direct current stimulation and transcranial magnetic stimulation of the cortex have shown that synchronized brain activity can enhance sleep slow waves (Marshall et al., 2006; Massimini et al., 2007), and several studies in humans and mice have found sleep-promoting effects of rocking and brief tones delivered at low frequencies (<u>≤</u> 1.5 Hz) (Bayer et al., 2011; Kompotis et al., 2019; Perrault et al., 2019; Tononi et al., 2010).

An alternative model, which we will refer to as the habituation model, proposes that habituation, a form of non-associative learning that is distinct from sensory adaptation or motor fatigue, plays a critical role in sleep induction by monotonous stimulation (Pavlov, 1927; Sokolov, 1963; Bohlin, 1971). The model suggests that habituation to repeated stimuli leads to reduced arousal and increased propensity for sleep through a common mechanism (Pavlov, 1927; Sokolov, 1963; Bohlin, 1971). Habituation is traditionally viewed as a process that allows organisms to ignore predictable, unimportant stimuli so they can focus on salient changes in the environment (Rankin et al., 2009; Thompson and Spencer, 1966). According to this view, once an organism has learned to ignore monotonous stimuli, they would no longer be effective at inducing sleep. However, recent findings suggest that habituation is more than simply learning to ignore unimportant stimuli and allows organisms to switch between alternative behaviors depending on environmental conditions (McDiarmid et al., 2019). Incorporating the more recent view of habituation, the model proposes that habituation allows organisms to choose sleep over wakefulness under monotonous stimulation conditions. The habituation and synchronization models are not necessarily mutually exclusive, and they may apply to varying degrees depending on the stimulus conditions.

Mechanosensory stimuli are processed by the auditory, vestibular (gravity sensing), somatosensory, and proprioceptive systems. The mammalian ear processes sound (vibration) and gravity in parallel auditory and vestibular systems, respectively. In the fly, the chordotonal organs in the antennae, wing bases and legs constitute major mechanosensory systems that mediate audition, gravity and wind sensing, and proprioception (Albert and Göpfert, 2015; Tuthill and Wilson, 2016). The antennal chordotonal organ of the fly, or Johnston’s organ, is analogous to the mammalian ear and consists of two main neuronal clusters responsible for the processing of sound vs. gravity/wind, respectively (Kamikouchi et al., 2009; Yorozu et al., 2009), whereas chordotonal organs in wing bases and legs mediate proprioception. A study in mice reported that rocking promotes sleep through the vestibular otolithic organs (Kompotis et al., 2019). However, several studies have demonstrated that repetitive acoustic stimuli can also enhance sleep slow waves in humans, presumably through the auditory system (Tononi et al., 2010; Bohlin, 1971). Together, these results suggest that mechanosensory stimuli can influence sleep through multiple sensory systems in mammals.

Sleep in *Drosophila melanogaster* exhibits several key features of human sleep (Hendricks et al., 2000; Joiner, 2016; Shaw et al., 2000). Due to the relative simplicity of its genome and the availability of sophisticated tools for precise spatial and temporal control of gene expression, *Drosophila* serves as a powerful model system for understanding the genetic and neural basis of sleep regulation (Allada et al., 2017; Artiushin and Sehgal, 2017; Cirelli, 2009; Kirszenblat and van Swinderen, 2019; Tomita et al., 2017). Discovering that mechanical stimulation promotes sleep in flies would allow efficient and detailed investigation of an intriguing phenomenon.

Here, we report that mechanosensory stimuli promote sleep in flies. Flies exhibited reduced sleep (‘negative rebound’) after vibration, which suggests vibration-induced sleep (VIS) leads to the accumulation of sleep credit. Flies exhibited reduced arousability during VIS relative to normal sleep, and heightened arousal through the circadian clock or elevated dopamine signaling counteracts sleep induction by vibration. We found that sleep latency decreases and sleep amount increases over successive blocks of vibration, suggesting that habituation contributes to VIS. Ablation of the antennae or chordotonal organs partially suppresses VIS but does not eliminate it, indicating that multiple sensory organs are involved. By presenting simple sinusoidal vibrations to three control strains, we found that vibrations ranging from 3 Hz to 200 Hz can induce sleep. The magnitude of VIS depended on the stimulus parameters and genetic background. Our data suggest habituation is a major contributor to VIS, but synchronization of brain activity may play a role under certain stimulus conditions.

## Results

### Gentle Mechanical Stimulation Promotes Sleep

To test whether gentle mechanical stimuli can promote sleep in *Drosophila*, we first placed *Drosophila* Activity Monitors (DAMs) on a shelf ∼40 cm above a multi-tube vortexer, such that a small amplitude vibration from the vortexer was coupled to the DAMs. After establishing a day of baseline sleep/wake behavior, we applied continuous vibration for a day to three control strains: *Canton-S* (*CS*), *iso31* (a commonly used *white*^1118^ control strain), and *CSx-iso31* (a derivative of *iso31*, in which the X chromosome is replaced by that of the *CS* strain). We found that daytime sleep in both males and females of all three strains was markedly increased during vibration (Figures 1A and 1B). The effects of vibration on nighttime sleep were modest or absent, which may be due to high levels of baseline sleep. Notably, sleep during vibration exhibited a normal decrease toward the end of the light period (Figure 1A), demonstrating that the circadian arousal signal modulates the effects of vibration on sleep and that the flies did not have difficulty moving during vibration. Video recording of their behavior revealed that flies initially responded to vibration with increased locomotor activity (Supplemental Movie 1), further confirming that vibration did not cause paralysis or difficulty in locomotion. Video recording also showed that flies gradually became inactive during vibration and that they did not engage in small movements such as eating or grooming that are not detected by the DAM System.

**Figure 1.**
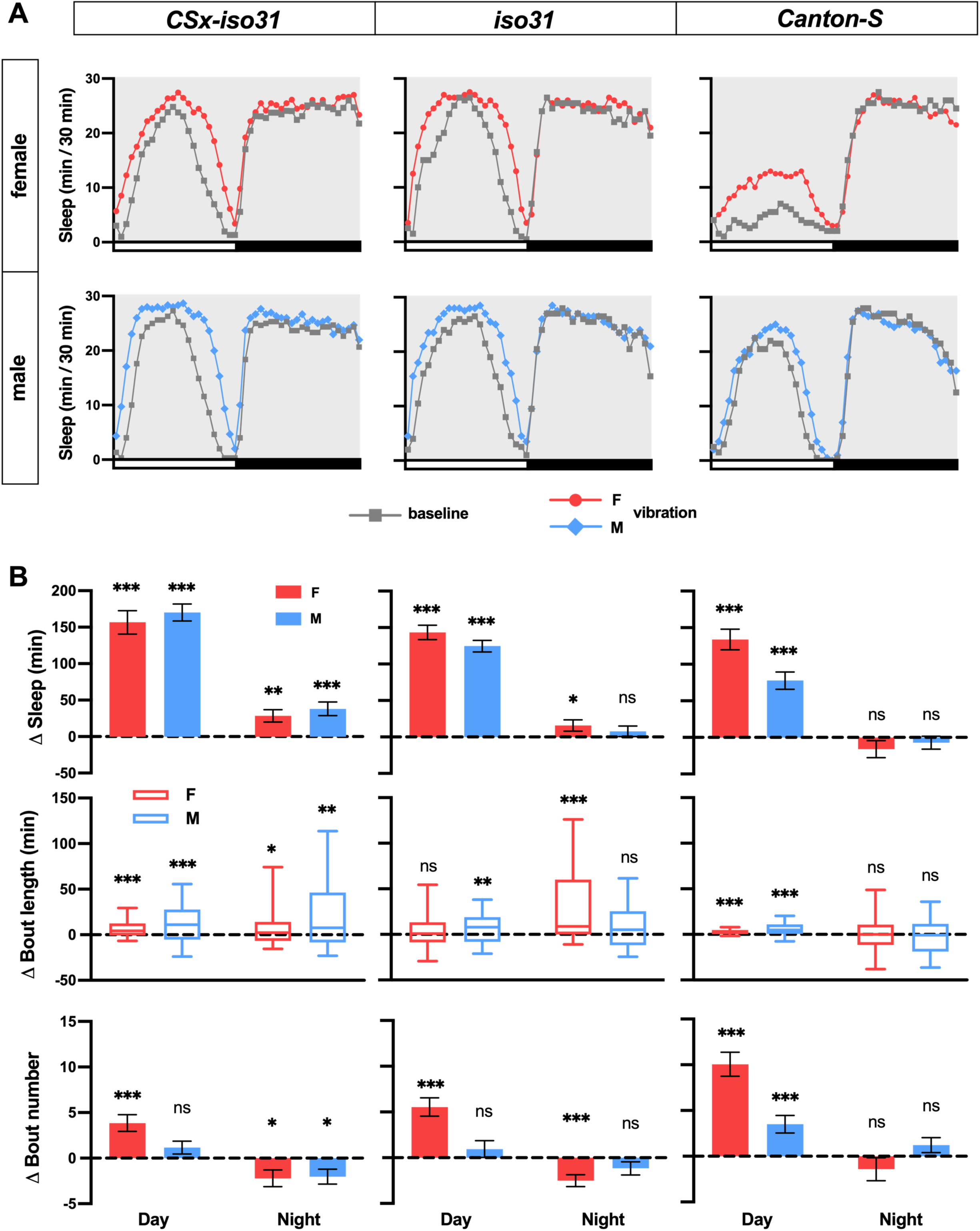
Vibration promotes sleep in flies. **(A)** Sleep profiles of *CSx-iso31* (left), *iso31* (middle), and *CS* (right) females (top) and males (bottom) exposed to vibration for 24 h starting at ZT0. n = 61-95. **(B)** Daytime and nighttime changes (vibration day vs. baseline day) in sleep amount (top), sleep bout length (middle), and sleep bouts (bottom) of flies shown in A. Error bars indicate s.e.m. For bout duration, the line inside the box indicates the median, and the whiskers indicate 10% and 90% percentiles. ns: not significant, *p < 0.05, **p < 0.01 and ***p < 0.001, paired Student’s t test (Δsleep and Δbout number) or Wilcoxon matched-pairs signed rank test (Δbout duration) with Bonferroni correction.

To examine the changes in sleep architecture during vibration, we examined sleep bout duration and bout number. The substantial increase in daytime sleep during vibration in all three strains was due to increased bout duration and/or number, whereas the modest nighttime sleep gain in *CSx-iso31* flies and *iso31* females was due to a combination of an increase in sleep bout duration and a decrease in sleep bout number (Figure 1B). These results suggest that vibration promotes sleep by influencing both sleep initiation (as reflected in increased sleep bout number) and maintenance (as reflected in increased sleep bout duration) depending on the genetic background and time of day. Collectively, our data establish that vibration induces sleep in *Drosophila*.

### VIS Results in the Accrual of Sleep Credit and is Independent of Light and the Circadian Clock

If sleep during vibration functions as normal sleep, we expect it to lead to a reduction of homeostatic sleep drive and an increase in sleep credit. To examine whether increased sleep during vibration contributes to the accumulation of sleep credit, we subjected *CSx-iso31* females to 6 h of vibration in the first half of the day. The end of vibration occurred at the peak of mid-day siesta when any decrease in sleep would be readily detectable. We observed a significant decrease in sleep in the 6 h after vibration, or negative rebound, following a substantial increase in sleep during the 6 h vibration (Figures 2A and 2B). These data show that sleep gained during vibration can contribute to sleep credit and lead to reduced sleep after vibration, suggesting that VIS can substitute for normal sleep.

**Figure 2.**
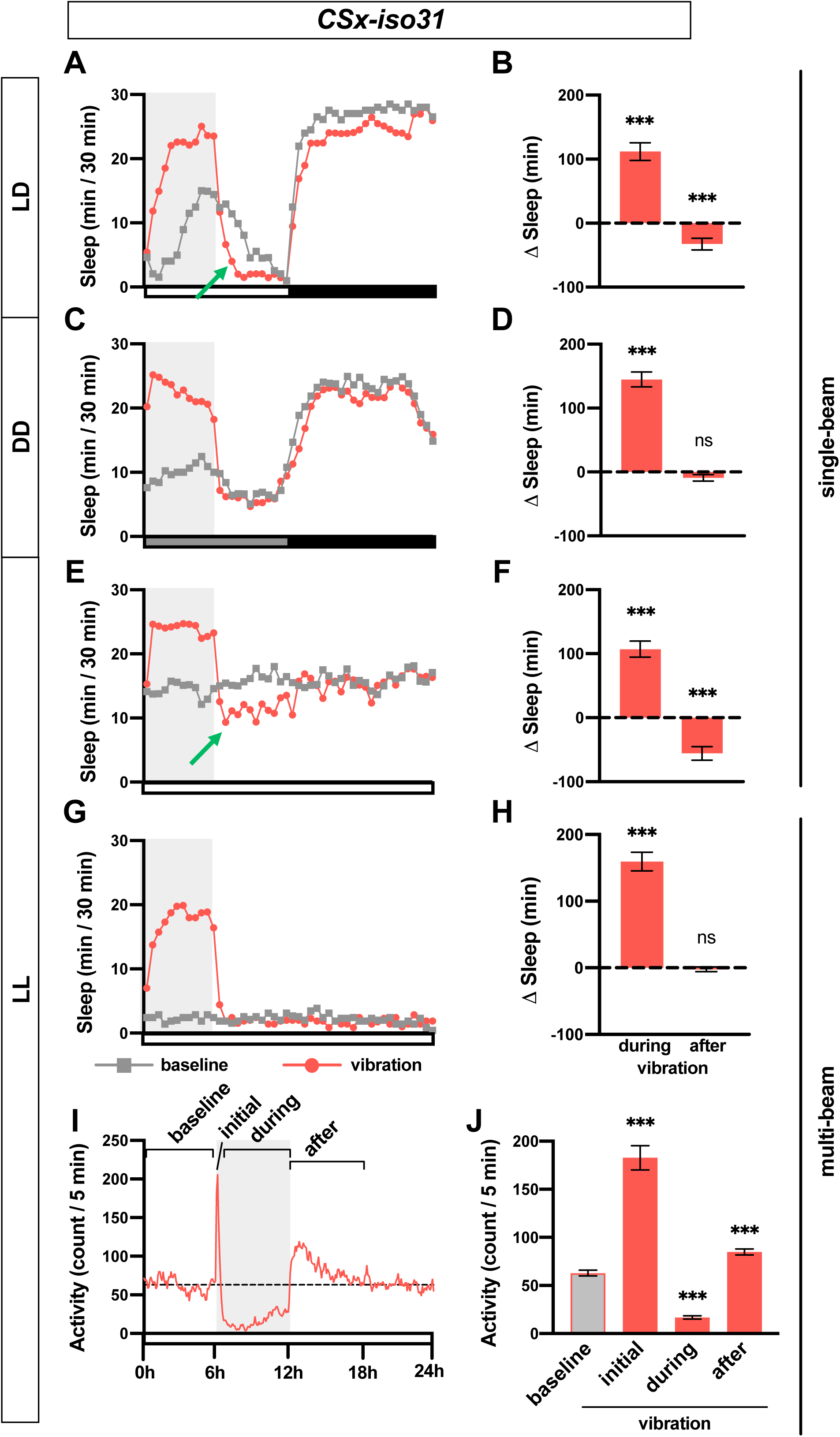
VIS results in the accrual of sleep credit and is independent of light and the circadian clock. **(A, C, E, G)** Sleep profiles of *CSx-iso31* females in LD **(A)**, DD **(C)**, and LL **(E, G)**, exposed to vibration for 6 h starting at Zeitgeber Time (ZT) 0 or Circadian Time (CT) 0. Single-beam monitors were used in A, C, and E, whereas multi-beam monitors were used in G. n = 30–75. Gray box indicates the 6 h period of vibration. Green arrows point to negative rebound. **(B, D, F, H)** Sleep change during or after 6 h vibration relative to a 6 h baseline period prior to vibration of flies shown in A, C, E, G, respectively. **(I)** Activity profile of flies shown in G. Both inter-beam and intra-beam movements in multi-beam monitors are included. **(J)** Average activity count during 6 h prior to vibration (baseline), first 5 min (initial) or 0.5-6 h (during) of vibration, and 6 h after vibration (after). Error bars indicate s.e.m. ns: not significant. *p < 0.05, **p < 0.01 and ***p < 0.001, paired Student’s t test with Bonferroni correction (B, D, F, H), repeated-measures ANOVA followed by Dunnett’s posthoc test relative to baseline (J).

Since our results showed a greater increase in sleep during daytime compared to nighttime under LD conditions (Figures 1A and 1B), we asked whether sleep increase by mechanical stimulation requires light or the circadian clock. To test this, we assayed sleep change during vibration in constant dark (DD) and constant light (LL) conditions. Flies exhibited clear VIS during the subjective day in DD (Figures 2C and 2D), demonstrating that light is dispensable for the phenomenon. In fact, the sleep increase during vibration was more pronounced in DD than in LD. However, flies did not exhibit negative rebound after vibration in DD, which suggests that homeostatic response to sleep credit is gated by light. In LL, where flies became arrhythmic, they also responded to vibration with increased sleep (Figures 2E and 2F), suggesting the circadian clock is not required for sleep induction by vibration. As in LD, flies exhibited negative rebound following 6 h vibration in LL, confirming that VIS contributes to the accrual of sleep credit. Our data demonstrate that vibration induces sleep independent of the circadian clock and light.

Sleep data presented thus far were obtained using the single-beam *Drosophila* Activity Monitoring (DAM) system, in which each fly is monitored by a single infrared detector. To determine whether the apparent reduction in activity during vibration was due to local movements that may not be detected by single-beam monitors such as eating or grooming, we employed multi-beam monitors containing 17 infrared beams. Multi-beam monitors allowed measurements of local (intra-beam) movements that occur within a single beam such as grooming as well as beam-to-beam (inter-beam) movements. As previously shown (Garbe et al., 2015), sleep measured using multi-beam monitors was markedly lower than that measured using single beam monitors (compare Figures 2E vs. 2G). The baseline sleep was extremely low, presumably because intra-beam movements over-represent activity by including not only local movements such as grooming but minor twitches that occur during sleep (Garbe et al., 2015). Importantly, we observed a profound sleep gain during vibration in LL using multi-beam monitors even when a very sensitive measure of activity was used (Figures 2G and 2H). Examination of activity counts (combined intra- and inter-beam counts) showed that flies initially showed increased activity in response to vibration but their activity decreased gradually to levels below the baseline level (Figures 2I and 2J). Activity levels returned to pre-vibration levels a few hours after vibration ended. Similar changes were observed when inter-beam and intra-beam activity were analyzed separately (Figure S1), confirming that vibration suppresses both local intra-beam movements and locomotion across the monitor tubes. We did not observe negative rebound after vibration using multi-beam monitors (Figures 2G and 2H), which is likely due low baseline sleep. However, flies were more active in the first 6 h after vibration compared to baseline (Figures 2I and 2J), suggesting that after excessive sleep during vibration, flies become more active even if reduced sleep is not detected due to a floor effect.

### Sensory Responsiveness to Light is Reduced During VIS

One of the defining characteristics of sleep is increased arousal threshold (Campbell and Tobler, 1984). To determine how vibration affects arousability, we compared the probability of sleeping flies to awaken in response to light during periods of vibration and no vibration. We performed the assay in LL, where flies sleep a moderate amount throughout the day, as it renders flies arrhythmic and thus eliminates the need to control for circadian fluctuations in arousability. We found that 1 min of extremely bright light (∼15,000 lux) superimposed on constant, moderate light (∼500 lux) can awaken 40-75% of sleeping flies within 2 min under baseline (no vibration) conditions (Figure 3A). We, therefore, used bright light of varying durations (1 sec, 15 sec, and 1 min) to measure sensory responsiveness in sleeping flies during vibration compared to no vibration. We found that vibration substantially reduced the responsiveness of flies to visual stimuli (Figure 3A). This was true for both males and females of two different wild-type strains at all stimulus durations except for 1-sec light stimulation of *iso31* males. Spontaneous awakening in the absence of light pulses was also reduced during vibration in *CSx-iso31* males and *iso31* females, but the greater effects of vibration in light pulse conditions indicate that sensory responsiveness is reduced by vibration. Whereas 1 min of bright light in the context of moderate light awakened only ∼10-30% of sleeping flies during vibration, 1 min of dark pulses were sufficient to awaken >75% of them (Figure S2A), showing that VIS can be reversed by salient changes in the visual environment. These results show that sleep during vibration is associated with reduced arousability relative to baseline sleep, which suggests that sleep during vibration is deeper than baseline sleep.

**Figure 3.**
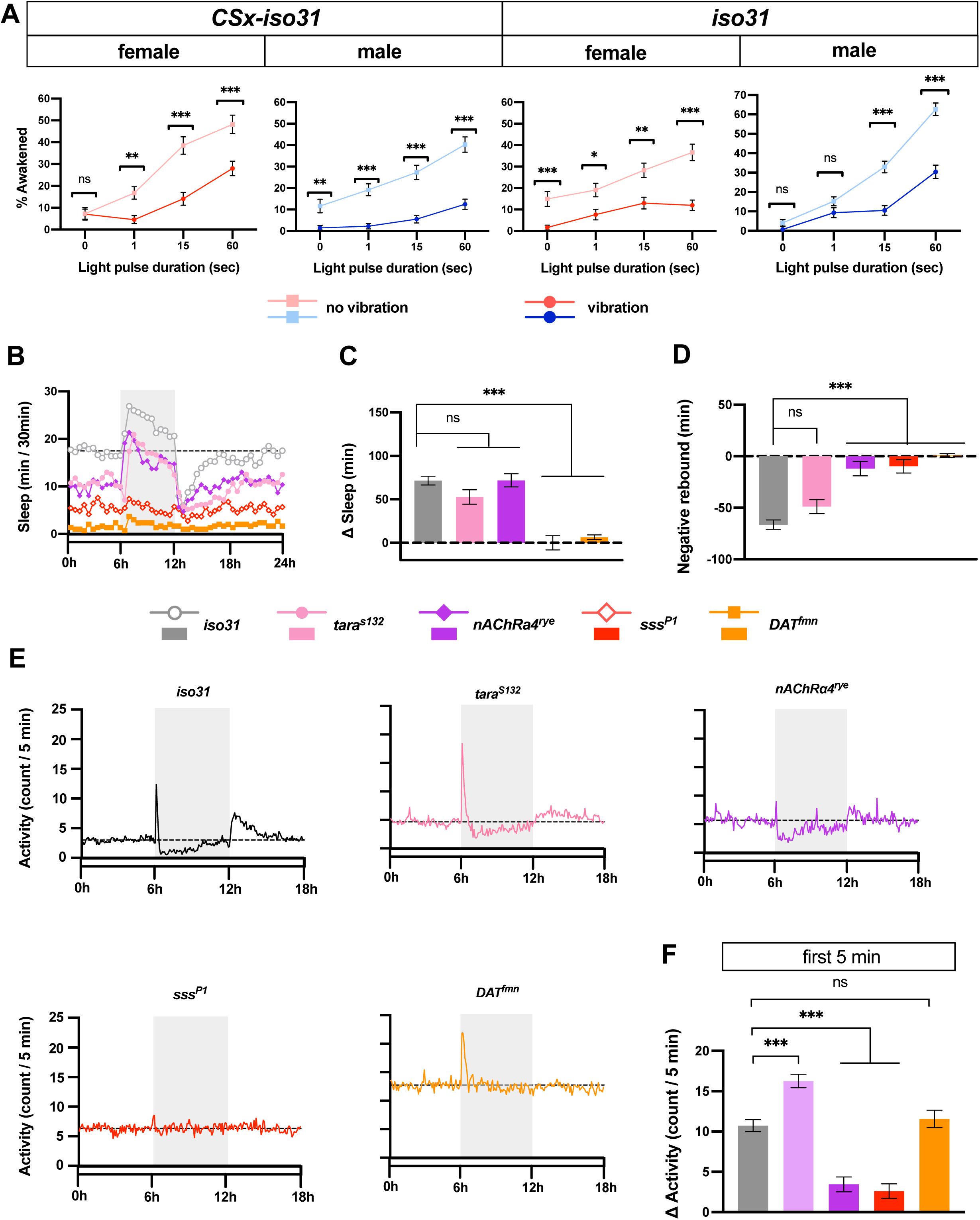
Sensory responsiveness to light is reduced during VIS and vibration has variable effects on short-sleeping mutants. **(A)** Percentage of sleeping flies that start moving within 2 min in response to bright light during periods of vibration or no vibration. *iso31* and *CSx-iso31* males and females were presented with bright light lasting 1 sec, 15 sec or 1 min. The “0 sec” data represent spontaneous awakening in the absence of light stimuli. n = 83-246 from 48 flies. **(B)** Sleep profile of control (*iso31*), *tara*^*S132*^, *nAChRa4*^*rye*^, *sss*^*P1*^ and *DAT*^*fmn*^ females exposed to 6 h vibration in LL. n = 37-128. **(C-D)** Amount of sleep change during 6 h periods during **(C)** and after **(D)** vibration compared to baseline (6 h prior to vibration) for flies shown in B. **(E)** Activity profile of flies shown in B. (**F**) Activity of flies shown in B during the first 5 minutes of vibration. Gray box indicates the 6 h period of vibration. Dotted lines indicate average baseline sleep (B) or activity (E). Error bars indicate s.e.m. ns: not significant. *p < 0.05, **p < 0.01 and ***p < 0.001, χ-square test with Bonferroni correction (A), Brown-Forsythe and Welch ANOVA followed by Dunnett’s posthoc test relative to controls (C-D, F).

### Vibration Has Variable Effects on Short-Sleeping Mutants

We next asked whether genetic mutations that affect baseline sleep also influence VIS. We applied vibration to several short-sleeping mutants: *sleepless* (*sss*) / *quiver, Dopamine transporter* (*DAT*), *taranis* (*tara*), and *nAChRα4* (Afonso et al., 2015; Koh et al., 2008; Kume et al., 2005; Shi et al., 2014). The mutant and control flies were vibrated for 6 h in LL, and sleep during vibration was compared to baseline sleep during the preceding 6 h. All the mutants exhibited reduced baseline sleep in LL (Figures 3B and S2B). *tara*^*s132*^ and *nAChRα4*^*redeye (rye)*^ mutants exhibited substantial VIS (Figures 3B and 3C), suggesting that the short-sleeping phenotype of some mutants can be rescued by mechanosensory stimulation. Whereas *tara*^*s132*^ and *nAChRα4*^*rye*^ mutants showed similar sleep gains during vibration, only *tara* ^*s132*^ mutants exhibited a significant negative rebound (Figures 3B and 3D). How much sleep credit accumulates during VIS likely depends on several factors such as the rate of dissipation of sleep drive during sleep and the rate of dissipation of sleep credit during wakefulness, and our data suggest that these rates differ between *tara*^s*132*^ and *nAChRα4*^*rye*^ mutants. In contrast to *tara*^*s132*^ and *nAChRα4*^*rye*^ mutants, *sss*^*P1*^ and *DAT*^*fmn*^ mutants exhibited little change in sleep during vibration (Figures 3B and 3C), confirming that the genetic background has a substantial impact on the effectiveness of vibration as a means of inducing sleep. Whereas *tara*^*s132*^ mutants showed a normal pattern of locomotor activity in response to vibration, i.e., an initial increase followed by a gradual decrease to a level below the baseline level, *nAChRα4*^*rye*^ mutants exhibited only a modest initial increase followed by a rapid decrease (Figures 3E, 3F, and S2C), which suggests distinct response kinetics in different genetic backgrounds. In contrast, *sss*^*P1*^ mutants did not show a noticeable change in locomotion during vibration, suggesting that the loss of *sss*^*P1*^ may impact sensory processing of vibration. Only control flies and *tara*^s132^ mutants exhibited increased activity post vibration (Figures 3E and S2D), consistent with the above result that they were the only ones to show significant negative rebound (Figure 3D). Interestingly, although *DAT*^*fmn*^ mutants exhibited increased locomotion that gradually decreased over time, the activity level did not fall below the baseline level (Figures 3E, 3F, and S2B). *DAT*^*fmn*^ mutants harbor a genetic lesion in the *dopamine transporter* gene, which is expected to cause increased dopamine signaling and heightened arousal (Kume et al., 2005). It appears that *DAT*^f*mn*^ mutants stay aroused during vibration despite normal sensory processing of vibration. These results suggest that dopaminergic arousal signals can counteract the sleep-promoting effects of vibration.

### Habituation Learning Leads to Improved Sleep Induction in Successive Blocks of Vibration

As discussed above, flies initially responded to vibration by increasing locomotor activity but their activity gradually decreased to levels below the baseline. Decreased response to repeated stimuli could be due to habituation, a form of non-associative learning. To test whether habituation to vibration alters sleep induction, we presented them with a series of 1 h vibration training blocks interspersed with 1 h rest periods. Habituation can be distinguished from sensory adaptation or motor fatigue by stimulus specificity, namely a stimulus similar to, but distinct from, the habituated stimulus can restore response strength to pre-habituation levels (Rankin et al., 2009; Thompson and Spencer, 1966). To assess stimulus specificity, we compared continuous vibration to intermittent (2 min on, 2 min off) vibration. Flies were exposed to either four blocks of continuous vibration followed by one block of intermittent vibration or four blocks of intermittent vibration followed by one block of continuous vibration. Both continuous and intermittent vibration were able to induce sleep, and the sleep-promoting effects increased over the four blocks (Figures 4A and 4B). Continuous vibration was more effective in inducing sleep than intermittent vibration in the first block as evidenced by greater sleep gain and shorter sleep latency, but the difference in sleep latency disappeared by the fourth block (Figures 4B and 4C).

**Figure 4.**
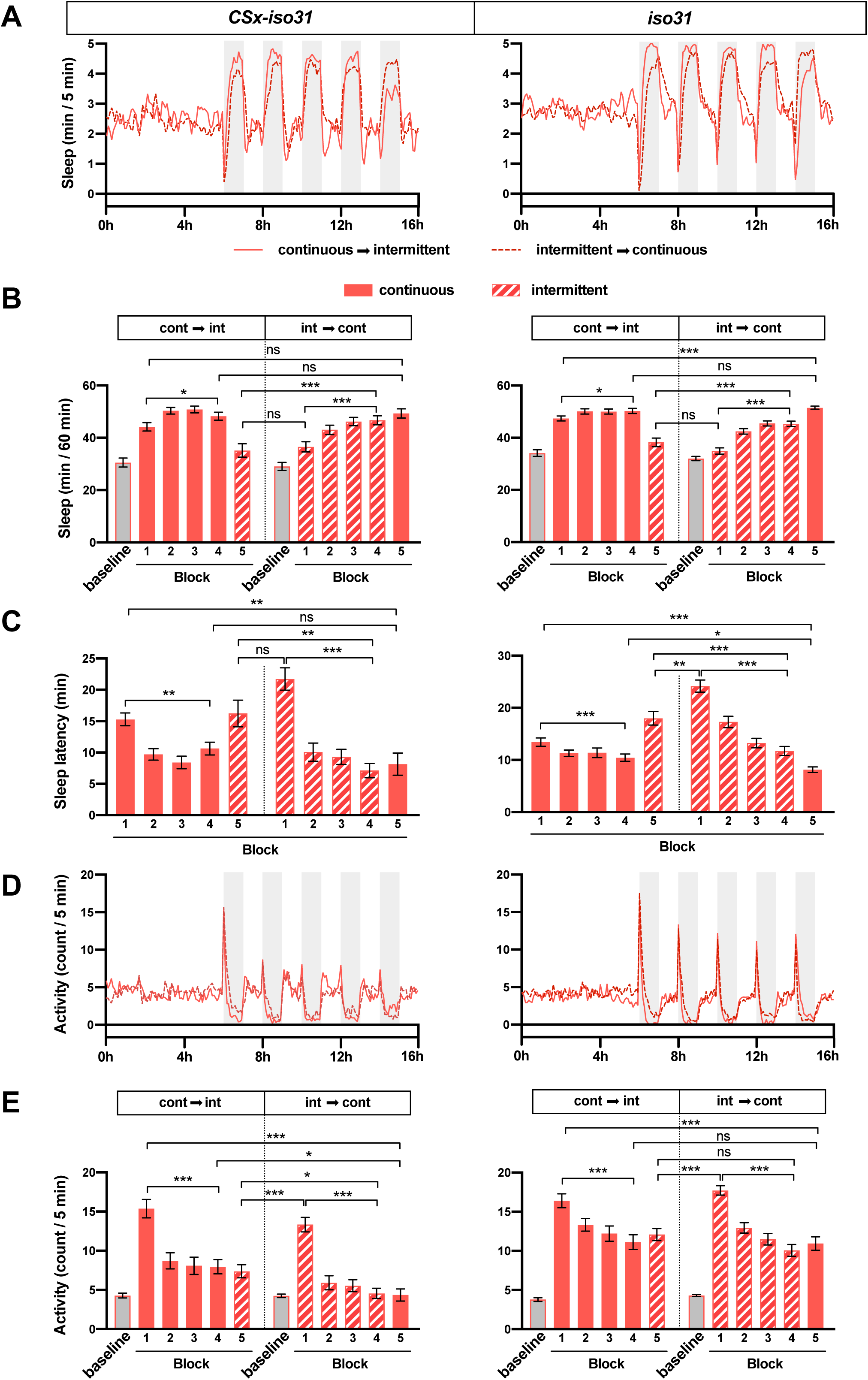
Habituation leads to decreased sleep latency in successive blocks of vibration. **(A)** Sleep profile of *CSx-iso31* and *iso31* females exposed to a series of 1 h blocks of vibration separated by 1 h of no vibration. Either 4 blocks of continuous vibration preceded 1 block of intermittent (2 min on, 2 min off) vibration (continuous → intermittent), or 4 blocks of intermittent vibration preceded 1 block of continuous vibration (intermittent → continuous). Gray boxes indicate the 1 h periods of vibration. **(B)** Sleep amount during the 6 h period prior to the first vibration block (baseline) and during blocks of 1 h vibration (block 1-5) for flies shown in A. Solid and striped bars represent continuous and intermittent stimulation, respectively. **(C)** Sleep latency relative to the onset of vibration in each 1 h block of vibration for flies shown in A. **(D)** Activity profile in 5 min bins of flies shown in A. **(E)** Average activity during the initial 5 min of vibration in each 1 h vibration block. Error bars indicate s.e.m. ns: not significant. *p < 0.05, **p < 0.01 and ***p < 0.001, unpaired Student’s t test with Bonferroni correction to compare a continuous vibration block vs. an intermittent vibration block, repeated measures ANOVAs followed by Dunnett’s posthoc test relative controls to compare blocks 1 vs. 4 within each experiment (B, C, E).

Switching from continuous to intermittent vibration and vice versa allowed us to examine stimulus specificity. Sleep in flies switched from continuous to intermittent vibration on the fifth block was more similar to sleep in flies experiencing intermittent vibration for the first time than those experiencing it for the fourth time (Figures 4B and 4C). This finding demonstrates stimulus specificity in that learning to fall asleep in response to continuous vibration did not generalize to intermittent vibration even though the stimuli are essentially identical except for the 2 min gaps between pulses of vibration. Some stimulus generalization was observed, however, especially with respect to locomotor activity. Activity levels in the first 5 min of vibration in flies switched from continuous to intermittent vibration on the fifth block were more similar to those of flies experiencing intermittent vibration for the fourth time than for the first time (Figures 4D and 4E). In contrast to switching from continuous to intermittent vibration, switching from intermittent to continuous vibration did not show stimulus specificity, which may represent a ceiling effect since flies exposed to continuous vibration reached asymptotic levels of sleep amount and latency after the first hour of stimulation. Our finding of stimulus specificity confirms that sensory adaptation or motor fatigue does not play a major role in VIS. Overall, our data demonstrate an improvement in sleep induction over multiple vibration sessions and suggest that habituation and consequent reduction in arousal contribute to VIS.

### VIS is Mediated in Part by the Chordotonal Organs and the Antennae

Neurons located in the chordotonal organs of antennae, wing bases, and legs carry out mechanosensation in flies including audition, gravity/wind sensing, and proprioception (Tuthill and Wilson, 2016). To determine whether chordotonal neurons are responsible for vibration sensing, we genetically ablated chordotonal neurons using a pan-chordotonal neuron *nan-*GAL4 driver (Kim et al., 2003) and the cell death gene *head involution defective* (*hid*) (Zhou et al., 1997). We observed a significantly lower amount of sleep increase in flies with ablated chordotonal neurons compared to genetic controls (Figures 5A and 5B), which suggests chordotonal neurons is a major mediator of sleep induction by vibration. The small residual effects of vibration in the flies with ablated chordotonal neurons may reflect the contribution of other mechanosensory organs such as bristles. Both optogenetic and thermogenetic activation of chordotonal neurons using CsChrimson and dTrpA1, respectively, promoted sleep (Figures 5C-5E), suggesting that the activity of these neurons can contribute to sleep regulation.

**Figure 5.**
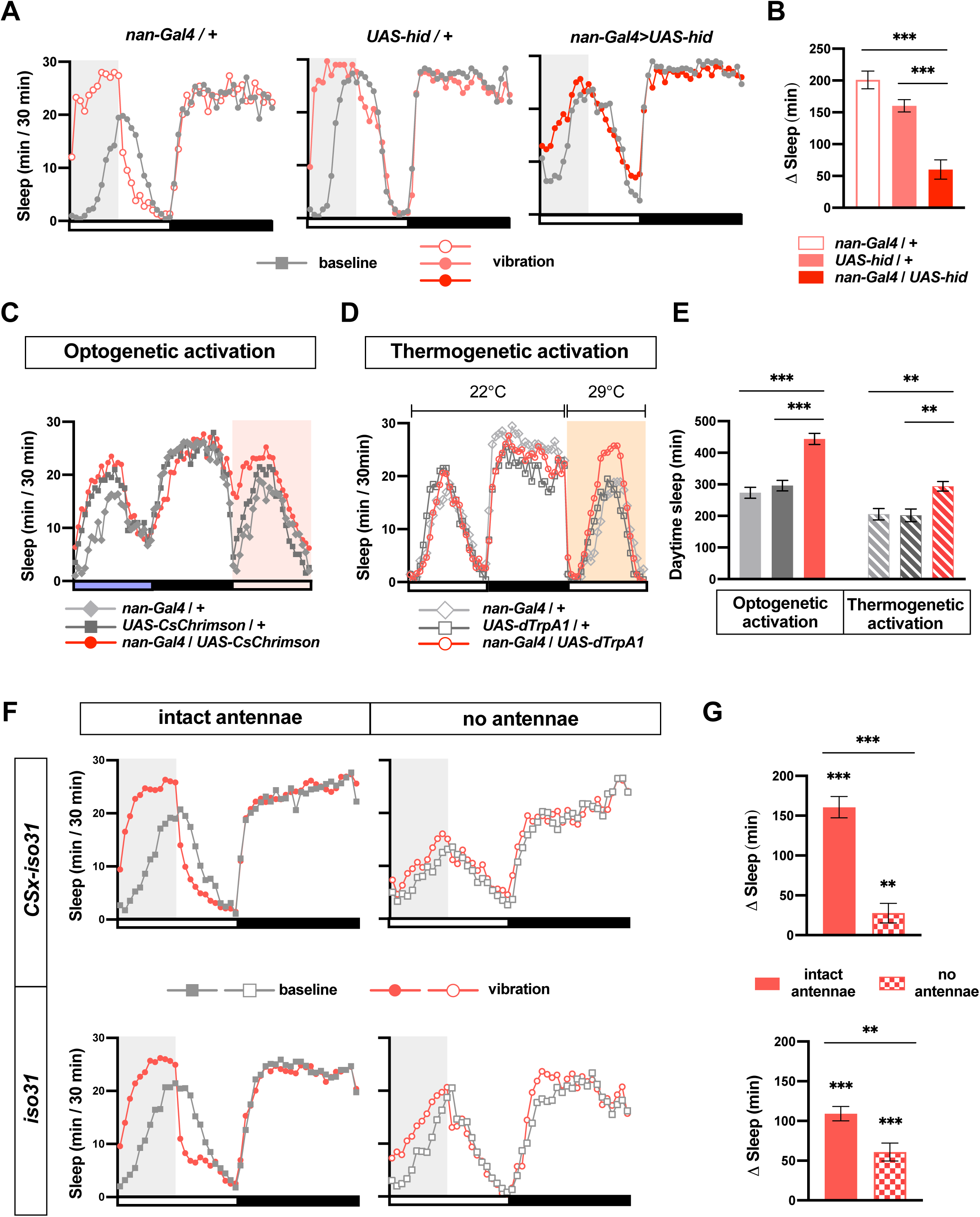
VIS is mediated in part by the chordotonal organs and the antennae. **(A)** Sleep profiles of nan*-Gal4/+, UAS-hid/+, nan-Gal4>UAS-hid* females. n = 24. **(B)** Sleep change during or after 6 h vibration relative to a 6 h period prior to vibration of flies shown in A. **(C)** Sleep profiles of female flies of indicated genotypes during baseline and optogenetic activation of chordotonal neurons. Red box indicates the period of exposure to red light. Blue light was used for entrainment to a 12h:12h LD cycle. n = 32-43. **(D)** Sleep profiles of female flies of indicated genotypes during baseline and during thermogenetic activation of chordotonal neurons. Orange box indicates the duration of exposure to high temperature. n = 29-41. **(E)** Total daytime sleep amount when flies are exposed to red light or high temperature for flies shown in C and D. **(F)** Sleep profiles of *CSx-iso31* (top) and *iso31* (bottom) females with intact antennae (left) or physically ablated antennae (right) exposed to vibration for 6 h starting at ZT0. n = 37-52. **(G)** Sleep change during or after 6 h vibration relative to a 6 h baseline period prior to vibration of flies shown in F. Gray box indicates the 6 hours period of vibration. Error bars indicate s.e.m. ns: not significant. *p < 0.05, **p < 0.01 and ***p < 0.001. One-way ANOVA followed by Sidak’s (B) or Dunnett’s (E) posthoc test; unpaired Student’s t test (to test whether sleep change is different in flies with intact antennae vs. flies without antennae) and paired Student’s t test (to test whether sleep change is different from 0) test with Bonferroni correction (G).

To determine whether the antennae, which contain a subset of chordotonal organs, mediate sleep induction by vibration, we physically ablated antennae of control flies and applied vibration. Both *CSx-iso31* and *iso31* flies exhibited significantly reduced VIS in the absence of their antennae compared to their peers with intact antennae (Figures 5F and 5G). Even without the antennae, both *CSx-iso31* and *iso31* flies showed a significant sleep increase during vibration relative to baseline levels. Overall, our results suggest that multiple sensory organs including chordotonal neurons in the antennae and elsewhere in the body convey mechanosensory information to the central sleep centers.

### Vibrations of a Wide Range of Frequencies Can Induce Sleep

Since vortexers produce a complex pattern of rotational and translational motions that cannot be easily manipulated parametrically, we built an audio loudspeaker-based system that allowed us to produce vertical translational motions with independently controlled frequency and amplitude. Our speaker system was built based on a design previously used to study circadian entrainment by vibration (Simoni et al., 2014) (Figure 6A). We started with a combination of 20 Hz and 200 Hz sinusoidal stimuli as similar frequencies were used in the circadian entrainment experiments. We found that 24 h vibration from the speaker system in LD had a profound effect on daytime sleep in both male and female *CS* flies (Figures 6B and S3). Vibration promoted sleep in *CSx-iso31* and *iso31* flies as well, although the effects were not as pronounced as in *CS* flies, perhaps due to higher baseline sleep during the day. As was the case with vibration generated by a vortexer, the effects of vibration on nighttime sleep were not as strong as those on daytime sleep. These data demonstrate that vibration generated by our speaker system can induce sleep in multiple control strains.

**Figure 6.**
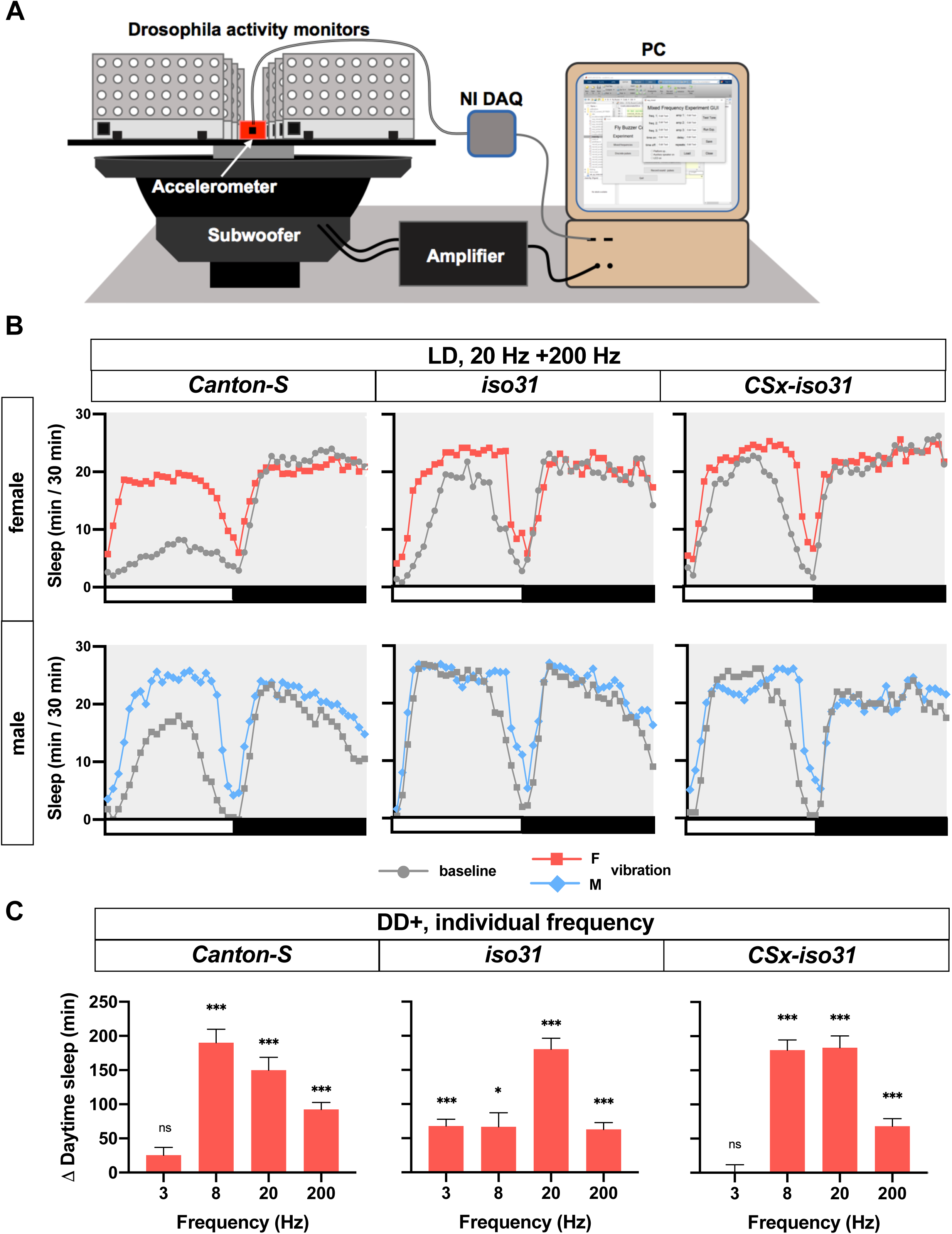
VIS depends on the stimulus frequency and genetic background. **(A)** Schematic representation of speaker system. Fly activity monitors were fastened to a platform, which was glued to the cone of a 15-in marine subwoofer. See Methods for additional details. **(B)** Sleep profiles of *CS, iso31, CSx-iso31* females (top) and males (bottom) exposed to a combination of 20 Hz and 200 Hz vibration for 24 hours in LD starting at ZT0. The amplitude of acceleration (half the difference between peak and trough) was 22.68 m/sec^2^. n = 43-108. **(C)** Daytime sleep change relative to baseline day of *CS, iso31*, and *CSx-iso31* females exposed to 24 hours vibration of 3 Hz, 8 Hz, 20 Hz, and 200 Hz. Sleep assay was performed in a “DD+” condition, i.e., darkness except for 5 min light pulses at ZT0 and ZT12. The amplitude of acceleration of 3, 8, 20, and 200 Hz vibration was 2.43, 15.78, 13.58, and 14.56 m/sec^2^, respectively. n = 51-64. Error bars indicate s.e.m. ns: not significant. *p < 0.05, **p < 0.01 and ***p < 0.001, paired Student’s t test with Bonferroni correction (B-C).

To determine the effects of vibration frequency on sleep, we applied vibrations at varying frequencies. Based on our data showing that VIS is more readily detectable in DD than in LD (Figure 2), we performed these experiments in a modified DD condition, in which it was dark except for 5 min light periods at ZT0 and ZT12. This lighting scheme, referred to as a “skeleton photoperiod” (Pittendrigh and Minis, 1964), allowed us to take advantage of greater vibration effects in darkness while minimizing the effects of variable circadian period lengths across individuals and genotypes. In addition to 20 Hz and 200 Hz vibrations administered separately, we included 3 Hz and 8 Hz vibrations. We used 3 Hz because it is close to the frequency range used in mammalian studies of rocking (Bayer et al., 2011; Kompotis et al., 2019; Perrault et al., 2019), and 8 Hz because a previous electrophysiological study found that the fly brain sometimes exhibits 7-10 Hz oscillations during sleep (Yap et al., 2017). Vibrations at all four frequencies were capable of inducing sleep, at least in some control strains (Figures 6C). Research in mice has found that within a range of 0.16 - 1.5 Hz, acceleration determines the magnitude of sleep induced by rocking (Kompotis et al., 2019). In our study, although the 8, 20, and 200 Hz stimuli were comparable in acceleration (see legend for Figure 6C), they produced different amounts of sleep gain. For example, 200 Hz vibration was less efficient at inducing sleep than 20 Hz vibration in all three strains, while vibration at 8 Hz had different effects depending on the genetic background (Figure 6C). Due to the limitations of the speaker system, the 3 Hz vibration we used had a much lower amplitude of acceleration than the vibrations at other frequencies. Overall, our data show that a variety of vibratory stimuli can induce sleep, and the magnitude of sleep gain is a function of the vibration frequency and the genetic background.

## Discussion

Our data establish that as in humans and mice, gentle mechanosensory stimulation can promote sleep in *Drosophila*. Flies showed increased activity and decreased sleep around light-dark transitions during 24 h vibration, indicating that they did not experience difficulty in locomotion and that the circadian arousal signal can counteract the sleep-promoting effects of vibration. Our observations that flies initially reacted to vibration with vigorous locomotion and that they can be awakened with salient changes in the visual environment also confirm that VIS is unrelated to other types of suppressed locomotion such as that induced by wind or fear in flies (Gibson et al., 2015; Yorozu et al., 2009) or tonic immobility in birds (Gallup, 1977). Previous studies have suggested that flies transition between lighter and deeper sleep stages during extended sleep bouts (Yap et al., 2017). Our results demonstrate that sleep during vibration is associated with reduced arousability, which suggests sleep during vibration may be deeper than sleep during periods of no vibration. Moreover, flies exhibited reduced sleep after vibration, suggesting that excess sleep during vibration contributed to the accrual of sleep credit. Thus, VIS functions similarly to normal sleep in terms of its effect on sleep drive. An important future goal would be to determine whether VIS can provide other functions of sleep such as improved memory and longevity.

The habituation model of how sensory stimulation promotes sleep proposes that habituation to repetitive, unimportant stimuli leads to reduced arousal (Pavlov, 1927; Sokolov, 1963; Bohlin, 1971). According to the model, sensory inputs would result in increased sleep if they can decrease arousal to a level below the baseline level, independent of the stimulus frequency. Our result that vibration of a wide range of frequency (3-200 Hz) can induce sleep supports the habituation model. Our findings that flies fall asleep faster and stay asleep longer over successive blocks of vibration and that this improvement does not generalize from continuous vibration to intermittent vibration further support the habituation model. Moreover, *DAT*^fmn^ mutants, which exhibit increased arousal, are resistant to the effects of vibration, consistent with the view that reduced arousal is essential for sleep induction by sensory stimulation. We found that sleep amount and latency show stimulus specificity while activity count does not, which resembles several studies demonstrating that different response components exhibit distinct habituation kinetics and are controlled by distinct molecules and neural circuits in zebrafish and *C. elegans* (Flavell et al., 2013; McDiarmid et al., 2019; Randlett et al., 2019). These findings led to a recent proposal that habituation is more than simply learning to ignore and that it allows organisms to switch between alternative behaviors depending on the context (McDiarmid et al., 2019). Falling asleep when there is repetitive mechanical stimulation may provide as yet unidentified adaptive advantage. It is possible that monotonous gentle vibration signals a relatively safe environment for sleep.

An alternative model of the sleep-promoting effects of sensory stimulation is that sensory inputs can synchronize cortical activity and boost sleep slow waves (Bellesi et al., 2014; Perrault et al., 2019). Despite a recent study reporting ∼1 Hz oscillations in the sleep-regulatory neurons in the central complex (Raccuglia et al., 2019), it is unknown whether there are brain-wide oscillations within the delta frequency range during sleep in *Drosophila*. It is also unclear whether 200 Hz vibration can entrain cortical activity, and whether synchronized activity at such a high frequency can enhance sleep. However, the previous observation that the fly brain exhibits 7-10 Hz oscillations during some periods of sleep (Yap et al., 2017) raises the possibility that 8 Hz stimulation may promote sleep in part through the synchronization mechanism. Overall, our data are consistent with the habituation model, but the synchronization model may also apply under certain stimulus conditions.

We found that a variety of vibratory stimuli were capable of inducing sleep in flies. Compared to the wide range of vibration frequency used in our studies, human and mouse studies investigating the effects of rocking on adult sleep used a narrow range of frequencies (0.16 - 1.5 Hz) (Bayer et al., 2011; Kompotis et al., 2019; Perrault et al., 2019). Nevertheless, they found variable results in terms of the specific sleep parameters that were affected (e.g., sleep duration and sleep latency). Seemingly small differences in stimulus parameters and types of rocking motion may influence the effectiveness of mechanosensory stimulation for promoting sleep in humans. In addition, we observed that the effects of vibration on sleep were dependent on the genetic background. Even among wild-type control strains, the effects of vibration of the same frequency and amplitude differed, suggesting the phenomenon is easily modified by genetic variations. Consistent with this interpretation, various short-sleeping mutants exhibited differential responses to vibration. Whereas *tara*^*s132*^ and *nAChRα4*^*rye*^ mutants showed strong sleep gain in response to vibration, *sss*^*P1*^ and *DAT*^*fmn*^ showed little change in sleep. The differential effects may be due to distinct defects in either the central sleep-arousal circuits or peripheral sensory organs in various short-sleeping mutants. Further, we found that sleep induction by mechanical stimulation can be improved with training in flies. It may be essential to optimize the stimulus parameters for each individual over multiple sessions when applying mechanosensory stimulation to treat human sleep disorders.

A recent study employing rhythmic horizontal movements at 0.25 - 1.5 Hz found that the vestibular otolithic organs mediate the effects of rocking on sleep in mice (Kompotis et al., 2019). We find that multiple sensory organs including chordotonal organs in the antennae and the rest of body are involved in sleep induction by vibration in *Drosophila*. Most studies of the effects of sensory stimulation on sleep have used mechanosensory stimuli such as rocking or acoustic stimulation (Bayer et al., 2011; Bohlin, 1971; Kompotis et al., 2019; Perrault et al., 2019; Tononi et al., 2010), and whether stimulation in other sensory modalities also influences sleep is an interesting and unresolved question. A few studies reported that olfactory stimulation can promote sleep in humans and rats (Sano et al., 1998; Goel et al., 2005), but another study found that olfactory stimulation was not very effective at promoting sleep in humans (Tononi et al., 2010). The variable results may be due to differences in the odorants and how they were administered. *Drosophila* studies, given the relative ease of high throughput analysis, may help address the question of the influence of olfactory and other non-mechanosensory stimuli on sleep. Regardless of whether non-mechanosensory stimulation also influences sleep, sleep induction by sensory stimulation in flies provides an important platform for the study of neural and molecular mechanisms of sleep regulation and further investigation may help us develop non-invasive treatments for patients with sleep disorders.

## Author Contributions

Conceptualization and Methodology: A.O.-C. and K.K. Investigation: A.O.-C., J.R.B., S.I., and A.C. Resources: P.D.M. and C.F.-Y. Writing – Original Draft: A.O.-C. and K.K. Writing – Review & Editing: P.D.M., J.R.B., S.I., A.C., and C.F.-Y. Funding Acquisition and Supervision: K.K. and C.F.-Y.

## Acknowledgments

We thank Drs. Amita Sehgal and Kazuhiko Kume and the Bloomington Stock Center for fly stocks, Dr. Bill Joiner for the SleepLab software, and Dr. Matthew Kayser and members of the Koh lab for helpful suggestions for improving the manuscript. This work was supported by grants from the National Institutes of Health (R01NS086887 to K.K and R01NS084835 to C.F.-Y.).

## Methods

### Fly stocks

Flies were raised on standard food containing molasses, cornmeal and yeast at 25°C under a 12 hours-12 hours light-dark cycle. *iso31* (*w*^*1118*^) and *Canton-S* (CS) lines were obtained from Amita Sehgal, and *CSx-iso31* was generated by replacing the X chromosome of *iso31* with the X chromosome of *CS*. Fly lines carrying *nan-Gal4* (#24903) (Kim et al., 2003), *UAS-hid* (#65403) (Zhou et al., 1997), *UAS-CsChrimson* (#55136) (Franconville et al., 2018), *UAS-dTrpA1* (#26263) (Hamada et al., 2008) were obtained from the Bloomington Drosophila Stock Center. *DAT*^*fmn*^ line was obtained from Kazuhiko Kume (Kume et al., 2005) and *nAChRα4*^*rye*^ line was obtained from Amita Sehgal (Shi et al., 2014). *sss*^*P1*^ (Koh et al., 2008) and *tara*^*s132*^ (Afonso et al., 2015) were described previously. Fly lines were outcrossed to *iso31* for at least five generations, except for *CS* and *CSx-iso31*.

### Sleep analysis

For sleep analysis, 3- to 5-day-old flies entrained to either 12 h:12 h light:dark (LD) cycle or constant light (LL) for at least 3 days. Approximately 16 males and 16 females were housed together until they were individually loaded into glass tubes containing 5% sucrose and 2% agar. In optogenetic experiments, 10mM all trans-Retinal (ATR, CAT#R2500 Sigma-Aldrich, St. Louis, Missouri) was added to the food. Experiments were performed at 25°C except where noted. Activity data (beam breaks) were collected in 1 min bins using *Drosophila* Activity Monitoring (DAM) System (Trikinetics, Waltham, MA) to measure sleep defined as a period of inactivity lasting at least 5 min (Huber et al., 2004). Single-beam monitors were used except where the use of multi-beam monitors was specifically noted. For multi-beam data, the “counts” setting was used to detect the combined local, intra-beam movements and inter-beam movements, whereas the “moves” setting was used to detect inter-beam movements only. Intra-beam movements were calculated by subtracting moves from counts. Sleep parameters were analyzed using a custom MATLAB-based software SleepLab (William Joiner).

### Generation of vibration stimuli

Activity monitors and recording arenas containing flies were placed on a shelf ∼40 cm above an analog multi-tube vortexer (Fisher Scientific, Pittsburgh, PA) or secured on the speaker system platform using screws. For the vortexer experiments, the intensity was set to 3, and the duration and timing of the mechanical stimulation was controlled via LC4 Light Controller (Trikinetics, Waltham, MA). For the speaker system experiments, a custom MATLAB GUI was used to generate audio signals of arbitrary frequency and amplitude. The Fly activity monitors were securely fastened to an acrylic platform that was glued to the cone of a 15-in marine subwoofer (PLPW15D, Pyle Audio, Brooklyn, NY). A PC delivered audio signals to the amplifier that powered the subwoofer, driving mechanical oscillations. The MATLAB GUI also collected acceleration data from the accelerometer via the NI DAQ. The amplitude of acceleration of each stimulus was measured using a triple axis accelerometer breakout (ADXL337, Sparkfun Electronics, Niwot, CO) mounted on speaker system platform.

### Video Recording

Flies were loaded into 7 mm x 16 mm x 4 mm wells containing 5% sucrose and 2% agar. Videos were recorded with a digital camera (Sony DCR-SX63) and edited using iMovie.

### Analysis of sensory responsiveness

Flies were exposed to extremely bright light (∼15,000 lux) of varying durations (1 sec, 15 sec and 1 min) after being entrained under constant moderately bright light (∼500 lux). Bright light stimuli were applied every 30 min during alternating periods of vibration (1 h) and no vibration (1 or 2 h) using LC4 Light Controller (Trikinetics, Waltham, MA). The train of light stimulus started 25 min after the onset of the first vibration period. Sensory responsiveness was measured as the percentage of flies that started moving within two minutes of the bright light stimulus out of those that were asleep at the time of the stimulus presentation. Similar calculations were performed on data 10 min prior to light stimuli to measure spontaneous awakening. A series of 1 min dark pulses were applied every hour during alternating periods of 1 h vibration and 2 h silence, starting 45 min after the onset of the first vibration. Excel (Microsoft, Redmond, WA) was used to determine the percentage of sleeping flies that awakened within 2 min of bright light or dark pulses.

### Optogenetic and thermogenetic activation

For optogenetic experiments, flies were fed normal food supplemented with 10mM ATR for 2 days prior to being loaded into activity monitor tubes containing 5% sucrose and 2% agar supplemented with ATR. A blue LED panel (465nm wavelength, HQRP, Harrison, NJ) was used to entrain flies to LD cycle and a red LED panel (630nm wavelength, HQRP, Harrison, NJ) was used to activate the CsChrimson channel. For thermogenetic experiments, flies were raised at 22°C, monitored for 1 day at 22°C to determine baseline sleep levels and 1 day at 29°C to activate the dTrpA1 channel.

### Antennae ablation

All three antennal segments of 1-to-2-day-old flies were physically removed using 3mm Vannas Spring Scissors (Fine Science Tools, Foster City, CA). After 3 days of recovery, flies were loaded into tubes for sleep analysis.

### Quantification and Statistical Analysis

Statistical tests were performed using GraphPad Prism 8. To determine whether changes in response to vibration were significantly different from 0, paired Student’s t tests with Bonferroni corrections were used. For comparison of pairs of groups such as antennae ablated vs. intact flies, unpaired Student’s t tests were performed, and for comparison of 3 or more groups, ANOVAs were performed. If the groups had unequal variances, t-tests for unequal variances or the Brown-Forsythe and Welsh version of ANOVAs were used. Following ANOVAs, Dunnett’s or Sidak’s posthoc tests were performed depending on the type and number of posthoc tests. To analyze sleep bout duration data, Wilcoxon matched-pairs signed rank tests with Bonferroni corrections were performed. For the analysis of arousability by light and dark pulses, χ-square tests were performed followed by Bonferroni correction for multiple tests.

## Supplemental Figure Titles and Legends

**Figure S1.**
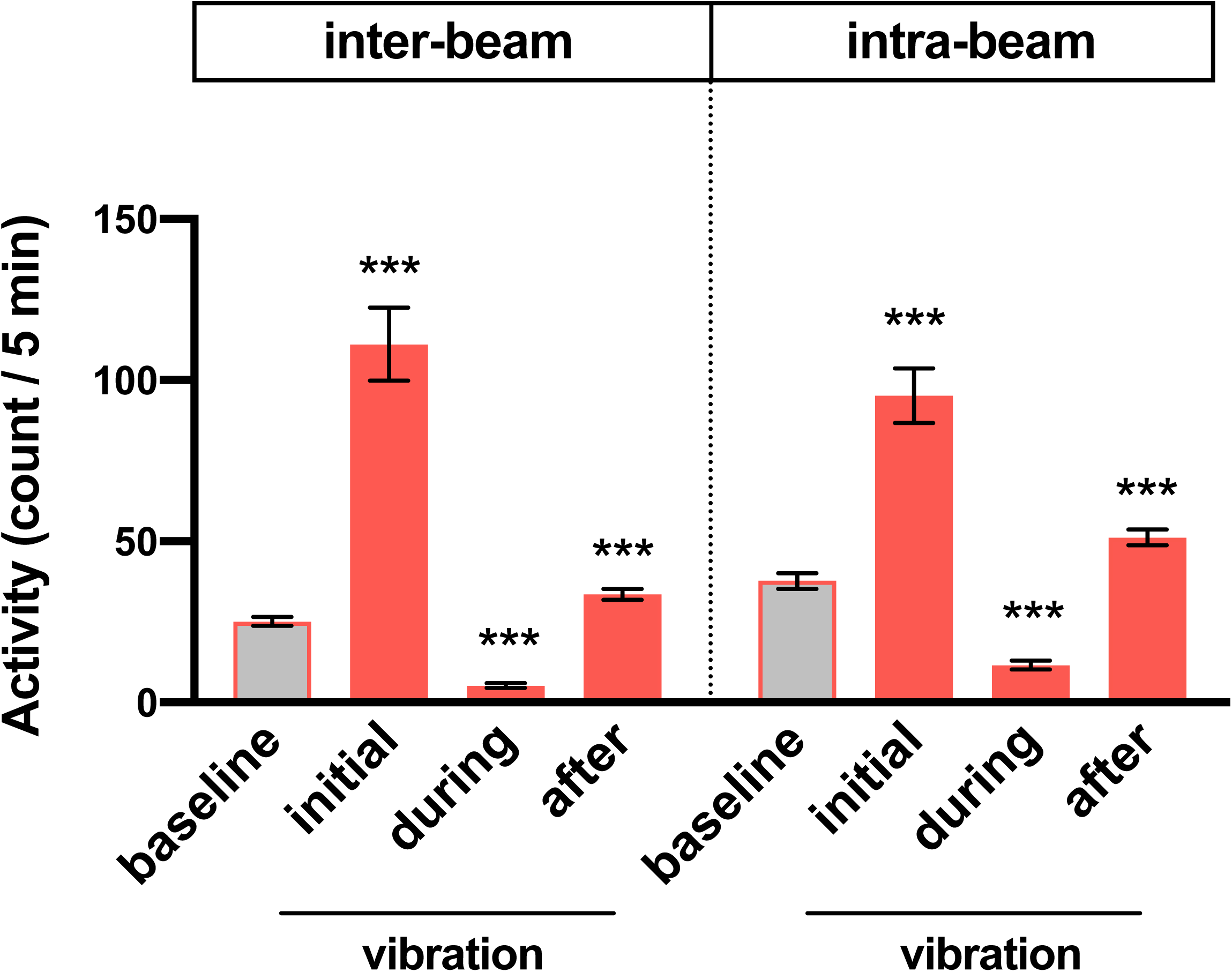
Vibration suppresses both local and non-local movements. Average inter-beam and intra-beam activity count during 6 h prior to vibration (baseline), first 5 min (initial) or 0.5-6 h (during) of vibration, and 6 h after vibration (after) for flies shown in Figure 2G. Error bars indicate s.e.m. ***p < 0.001, repeated-measures ANOVA followed by Dunnett’s posthoc test relative to baseline.

**Figure S2.**
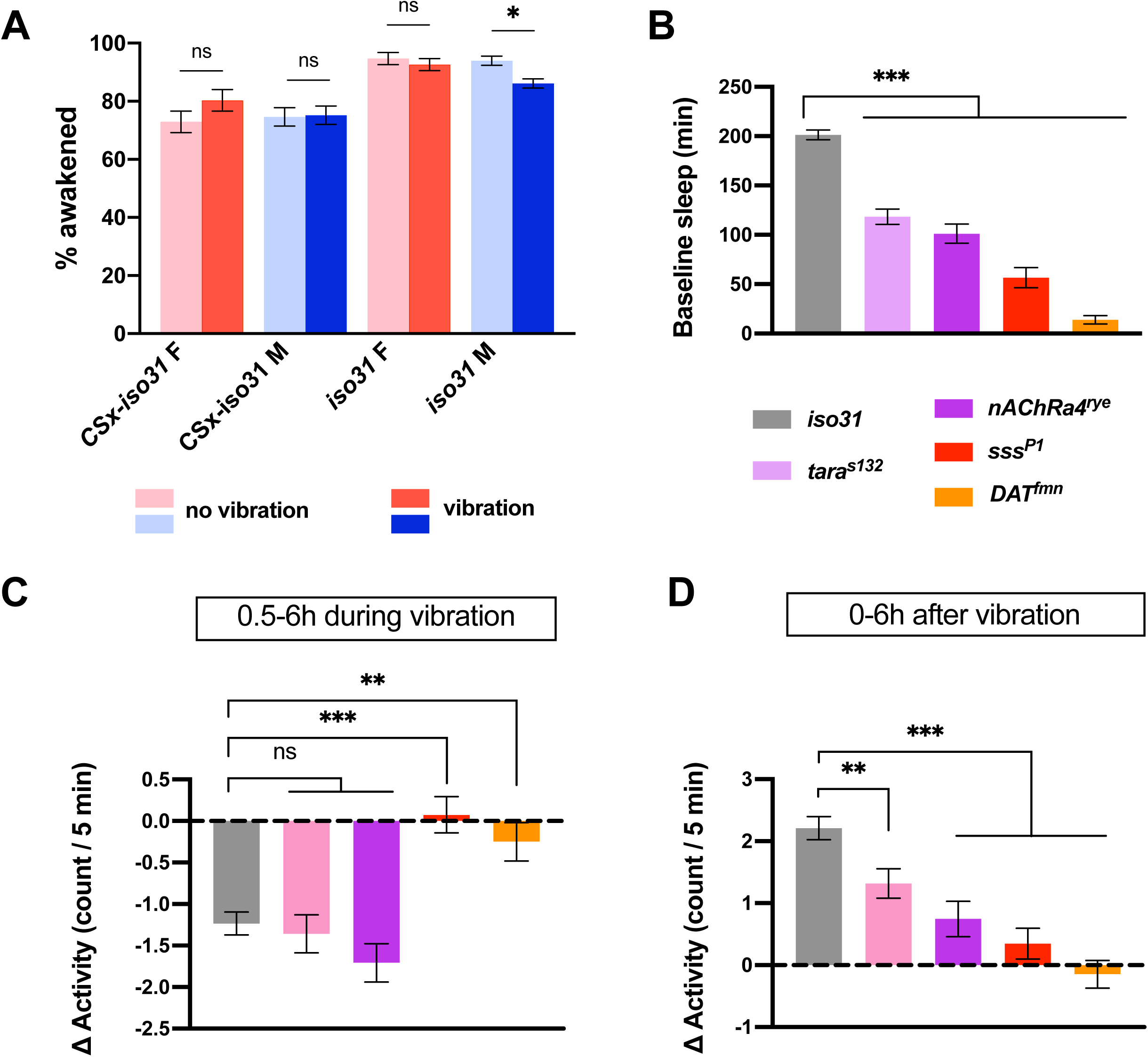
Vibration has variable effects on short-sleeping mutants. **(A)** Percentage of sleeping flies that start moving within 2 min in response to 1 min dark pulses during periods of vibration or no vibration. n = 114-231 from 61-64 flies. **(B-D)** Baseline sleep during 6 h prior to vibration (B), change in activity during vibration excluding the first 30 min (C) and 6 h after vibration (D) for females of the indicated genotypes shown in Figure 3B. Error bars indicate s.e.m. ns: not significant. **p < 0.01 and ***p < 0.001, χ-square test with Bonferroni correction (A), Brown-Forsythe and Welch ANOVA followed by Dunnett’s posthoc test relative to controls (B-D).

**Figure S3.**
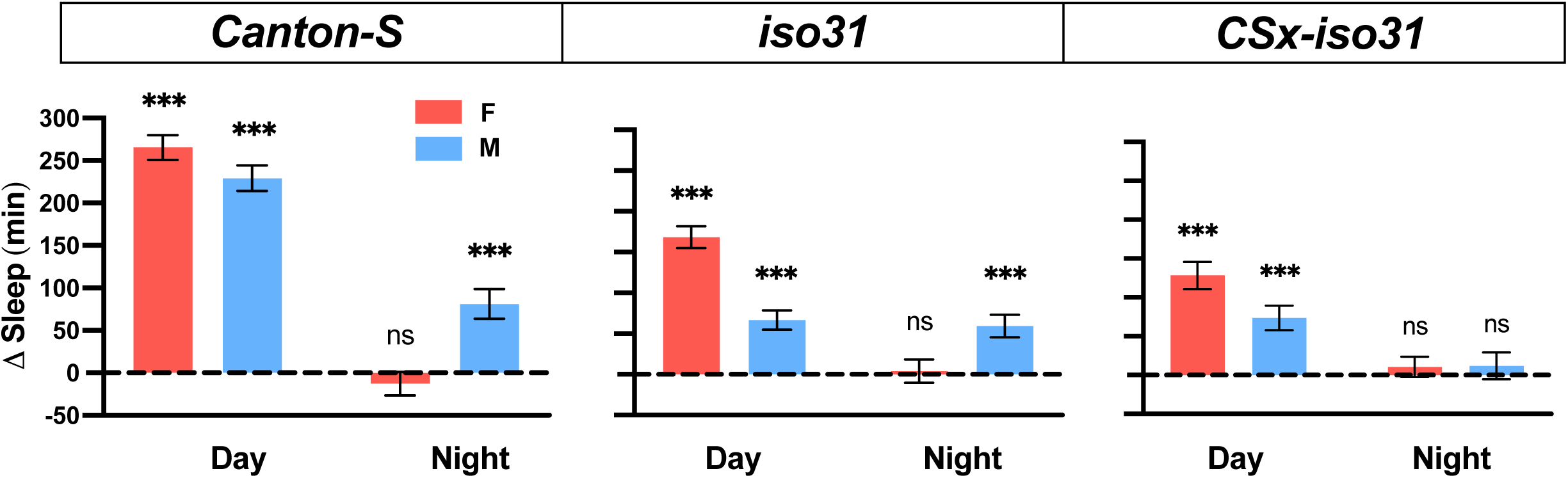
Sleep induction by vibration generated by a speaker-based system. Sleep change during daytime and nighttime relative to baseline of flies shown in Figure 6B. Error bars indicate s.e.m. ns: not significant. *p < 0.05, **p < 0.01 and ***p < 0.001, paired Student’s t test with Bonferroni correction.

**Supplemental Movie 1. Flies exhibit reduced activity during vibration.** Female *CS* flies were subjected to vibration for 1 h starting at ZT ∼3. Flies increased their activity at the start of vibration, but were mostly inactive 10 min later. The majority of them resumed their activity several min after the end of vibration.

